# Evaluation of FlowVR: a virtual reality game for improvement of depressive mood

**DOI:** 10.1101/451245

**Authors:** Lukas Bittner, Fariba Mostajeran, Frank Steinicke, Jürgen Gallinat, Simone Kühn

## Abstract

**Objective:** This study evaluated the efficacy of FlowVR, a virtual reality (VR) game designed to improve mood and reduce feelings of depression. The aim is to contribute to the question of whether and how VR could be used for depression therapy, as research in this area is quite rare.

**Method:** 18 healthy participants (9 female; *M*_age_ = 25.9) underwent three conditions, playing FlowVR in VR with a head-mounted display, playing FlowVR on a tablet or reading a text on a tablet. For each condition, they were tested on a separate day at the same time of day within a two-week period. Before and after every condition participants completed the Becks Depression Inventory II (BDI-II), the state part of the State-Trait-Anxiety-Depression-Inventory (STADI(S)) and the Positive Affect Negative Affect Schedule-Expanded Form (PANAS-X).

**Results:** While the participants showed only a reduction in acute anxiety in the control and the tablet conditions, they showed improved affectivity in all variables measured in the VR condition. In addition, VR had significantly better results than the control condition in improving positive affectivity, negative affectivity and acute feelings of depression. Using a less conservative statistical approach, these significant differences could also be found between the tablet and the VR condition. There were no significant differences between the tablet and the control condition.

**Conclusion:** The results indicate that due to its immersive nature, VR can be used effectively to improve mood and temporarily reduce feelings of depression. Long-term effects of FlowVR on participants with depression must be investigated in consecutive research.

## Introduction

According to World Health Organization, depression is the world’s leading cause of disability and strongly contributes to the global burden of disease ^1^. Since, as reported, only about one half of the 300 million people currently suffering from depression receive a treatment, innovative approaches to make therapy more efficient and accessible to the general public are needed. Virtual reality (VR) has already successfully been utilized for various therapy forms such as exposure therapy, in which patients can face their fears in a virtual environment (VE) in a safe and secure way. For example, soldiers who experienced traumatic events during the Vietnam War were exposed to their adverse experiences in VR and thus managed to significantly reduce their symptoms of anxiety and posttraumatic stress (Rothbaum, Hodges, & Kooper, 1997; Rothbaum, Hodges, Ready, Graap, & Alarcon, 2001). Many other disorders like schizophrenia, substance-related disorders and eating disorders were treated successfully as well (Freeman et al., 2017). However, Freeman and colleagues (Freeman et al., 2017) were only able to identify two studies regarding VR depression therapy when systematically reviewing the previous studies on VR therapy. Additionally, according to our research, no other papers on the topic have been published since then.

The first study was conducted by Falconer et al. (Falconer et al., 2016), who have been able to reduce depression in patients by increasing their level of self-compassion. This was accomplished by recording a VR session where the participant had to behave compassionately towards a virtual child. The participant was then embodied into that virtual child and received his own recorded compassion at the end. Secondly, Shah et al. (Shah et al., 2015) developed a VR stress management program where a combination of psychoeducation and VR-based relaxation practices significantly decreased levels of depression and anxiety. Additionally, Banos et al. (Baños et al., 2011) implemented a cognitive-behavioural therapy (CBT) program in their VR-world EMMA. They found statistically significantly better results in relaxation and depression compared to a traditional CBT program. Furthermore, a couple of guided meditation VR applications came up in the past years. Guided Meditation VR by Cubicle Ninjas^2^ or DEEP by Monoband-Play^3^ are designed to reduce the user’s stress, anxiety and discomfort levels. Research on these applications seems to be quite rare but substantiates positive effects on mental health (e.g. Perhakaran et al., 2016).

VR is defined as a “computer-generated simulation of a three-dimensional image or environment that can be interacted with in a seemingly real or physical way“^4^. The objective properties a device and the virtual environment (VE) offers to achieve this level of realism is described by the term ‘immersion’. The higher the level of immersion, the better the device simulates the VE and the interaction within it (Slater & Wilbur, 1997). When wearing the head-mounted display (HMD) of the HTC Vive^5^ for example, the users have a binocular view of the VE and their entire field of view is covered by it. Additionally, they can freely look around the VE similarly to being in a real world. On a tablet, in comparison, users have only a monocular view of the VE and they can visually perceive their real surrounding. Furthermore, the VE is not automatically adapting to their viewing direction. Consequently, a HMD can be categorized as VR device and is objectively more immersive than a tablet.

The subjective feeling of being in the virtual world is called the ‘sense of presence’ (Slater & Wilbur, 1997). When feeling present in the VE, the users are (re)acting similar to being in a real world (Slater & Wilbur, 1997). As the sense of presence increases, the more intensive and realistic emotional reactions, such as fear or joy, can be elicited by the VE (Regenbrecht, Schubert, & Friedmann, 1998). Therefore, the immersive nature of VR is the main reason of why VR can be efficiently used for psychotherapy (Strickland, 1996). Patients can be safely and securely exposed to their fears in the virtual world and still experience similar emotional reactions to those in reality (Regenbrecht et al., 1998). Likewise, if the patients feel present in the VE, their mood can also be influenced by the atmosphere and the experiences that they made within it (Riva et al., 2007). Consequently, if designed in the right way, a VR application could be particularly effective in improving patients’ moods.

Improving the mood of a patient can potentially have a long-lasting positive affect on her/his mental health. The so-called mood-congruence effect occurs, when people memorize or recall information in a distorted manner, influenced by their current mood (Wittchen & Hoyer, 2011). Being in a negative or positive mood increases the tendency to memorize and recall respectively negative or positive information. This effect is compatible with the Beck’s cognitive theory of depression (Beck, 1979), which states that depression is underlying dysfunctional basic assumptions and thought patterns (e.g. “I have to be perfect.”), that lead to a negative biased perception of reality. People in a depressed mood memorize and recall depressing memories more likely, thus develop and maintain negative cognitive schemata and a negative perspective on reality (Ingram, 1984). This repeated evocation of mood-congruent memories strengthens the preservation of depressive feelings. Consequently, inducing a positive mood can counteract the negative biased perception of reality. Once primed by a positive mood, negative experiences will less likely be remembered and positive experiences will more likely be recognized as positive. This has the potential to help patients to break the vicious circle of negative patterns of thought and depressive mood. Thus, inducing a positive mood could support a cognitive-behavioral therapy and foster the treatment of depression.

In the developed VR game, the player flies through a nature scene with the goal of making the landscape blossom. This can be achieved by the player flying near flowers, which are surrounded by a colored halo, to virtually pollinate them so that more flowers grow in the surrounding area. Visual and auditory elements associated with positive affect have been implemented and the task has been designed so that the players would make the experience of flow (i.e. being absorbed in the task), as can be seen in figure 1. The game is inspired by Thatgamecompany’s game Flower.

**Figure 1.**
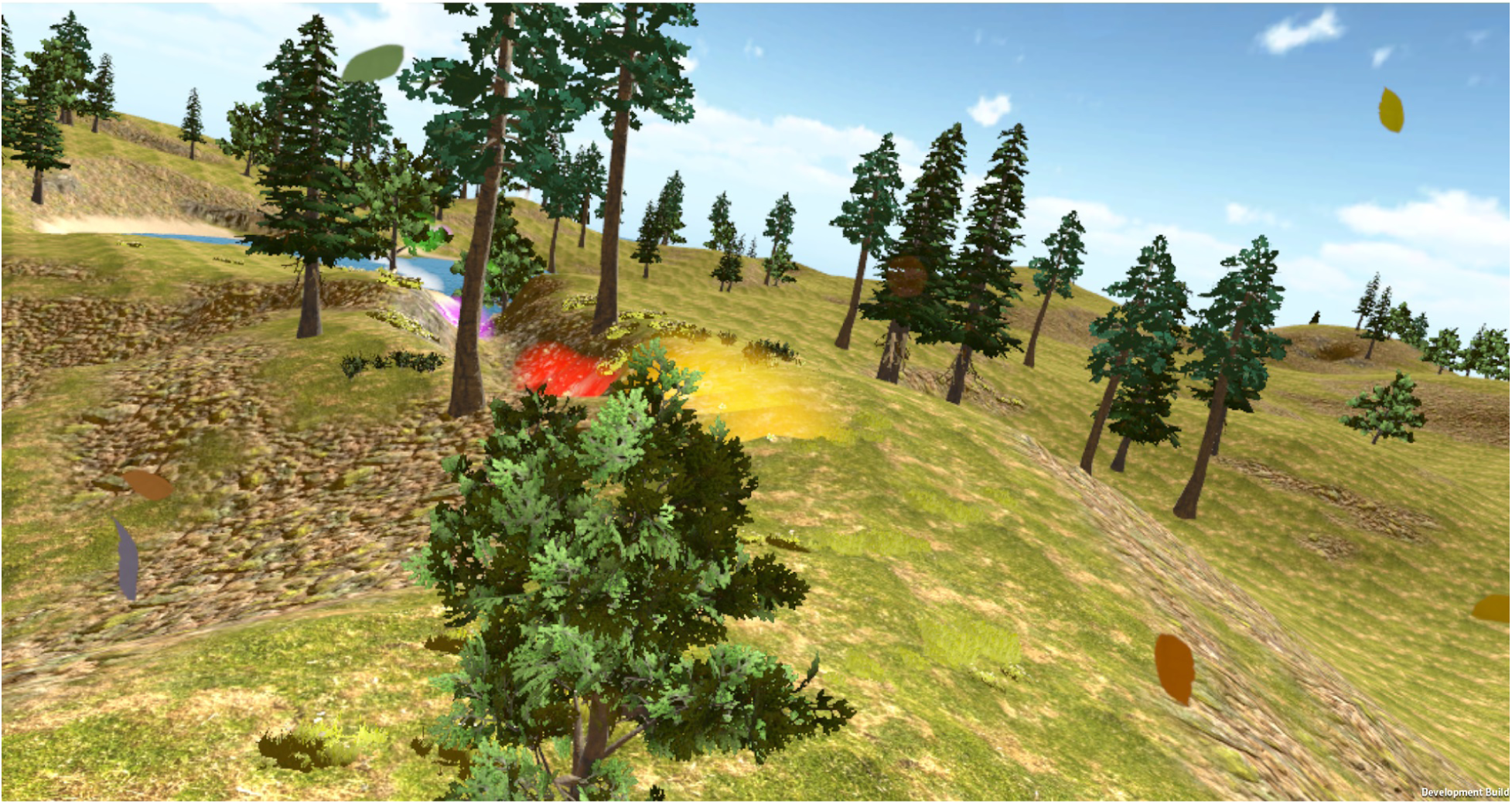
A screen-shot of the user’s view when flying above the virtual landscape. The colourful halos are indicating the positions of the next flowers to be ‘pollinated’. Leaves are circling around the user’s view.

It is well known that music can have a big impact on emotions and mood (Västfjäll, 2001). Even though there are individual differences in the perception of music, certain structural elements may have an impact on the emotional response. Especially mode, tempo, pitch and harmony of a song can be crucial elements for its emotional perception (Västfjäll, 2001). As a major mode is associated with a positive mood (Husain, Thompson, & Schellenberg, 2002; Västfjäll, 2001), the song implemented into the game was chosen to be in a major mode. Similarly, it was chosen to have higher pitches and simple harmonies (Rigg, 1964). As a fast tempo is associated with a higher arousal (Husain et al., 2002; Fernández-Sotos, Fernández-Caballero, & Latorre, 2016) but can be perceived as uneasy (Rigg, 1964), the song was preferred to be rather slow.

One experience that can have a particularly positive influence on mood, is called flow. To experience flow, one must be involved in a task, in which the level of difficulty matches the individual’s skills for managing the task (Csikszentniihalyi, 2014). Furthermore, a clear goal and immediate feedback have to be provided. The goal of the game is to make the landscape blossom and immediate feedback is provided by the new flowers growing immediately around the pollinated flower. As tasks which are too difficult could lead to overtaxing or anxiety (Csikszentniihalyi, 2014), the radius in which the flowers can be pollinated is set rather big to avoid that players will find the game too difficult (as indicated by the halo in figure 1). Additionally, this feeling of flow might be intensified by act of flying through the VE.

The response devices (controls) for VR and for tablet were designed with the same logic. In VR the player has to rotate his head to control the flight direction as he simply flies in the direction he is looking at. On the tablet, the user has to rotate the back of the tablet in the direction he wants to fly. To access the orientation of the tablet the gyroscope was used. If an arrow would stick out of the back of the tablet, it would indicate the current flying direction. Likewise, if an arrow would sick out of the VR’s HMD, it would indicate the current flying direction. The speed of flying was set low for both devices. These simple controls were chosen to make the conditions as comparable as possible and in order to avoid Simulator Sickness (SS), which is a well-known problem that can occur when it comes to the use of VR (Kolasinski, 1995). According to the Sensory Conflict Theory (Reason & Brand, 1975), it is caused by a conflict between the visual and the vestibular perception. Because there is no vestibular feedback when flying through the virtual world, there is a high chance that users will get sick.

The following hypotheses were formulated. (H1) Playing the game FlowVR, due to its positive elements, can improve mood and temporarily reduce depression compared to a control condition, i.e. reading a text on glider flights, regardless of the type of the medium on which it is played, i.e. HMD or tablet. (H2) The game in VR, due to its immersiveness, can provide a higher sense of presence compared to the game on a tablet and a control condition. (H3) Playing the game in VR, due to its higher sense of presence, can improve mood and temporarily reduce depression more than the game on a tablet.

## Method

### Participants

The study was submitted to the local ethical committee of the University of Hamburg. Candidates were recruited by a local job market website and an email distributor among the computer science students of the University of Hamburg. To apply for the study, they had to fill out the Mini International Neuropsychiatric Interview (MINI), Becks Depression Inventory II (BDI-II) and the trait part of the State-Trait-Anxiety-Depression-Inventory (STADI(T)) beforehand. The MINI is a short structured diagnostic interview for DSM-IV and ICD-10 psychiatric disorders (Hergueta, Baker, & Dunbar, 1998). The BDI-II assesses depression symptoms within the last two weeks and the STADI(T) assess depression and anxiety as a personality trait. For more details on the BDI-II and the STADI see section.

Applicants who had an acute Axis I disorder or acute alcohol or drug problems, according to the results of the MINI, were excluded. Furthermore, only participants with slightly elevated BDI scores (≥ 4) were tested, to make sure their level of depression could be improved after all. Out of 290 applicants, 43 fulfilled these criteria of which 19 took part in the study. One participant was later excluded, as he/she had been diagnosed with a mental disorder before and took antidepressants. Of the remaining 18 participants 9 were female and between 18 and 55 years old (mean=25.9, sd=10.5). None of those had ever been diagnosed with a mental disorder or took antidepressants. Two participants were taking a contraceptive pill and one participant took L-Thyroxin, a drug against hypothyroidism. Eight of the subjects were students in the computer science department. Four obtained class credit for their participation and the other 15 were paid 25 Euro. Six of the 18 participants had not worn a HMD before. Seven participants almost never played 3D video games, while four played on a monthly basis, three on a weekly basis and four on a daily basis. Six wore glasses or contact lenses and one subject had a red-green color blindness. The color blindness might have had reduced the positive effect of the green colors in the game. However, since this affects both the tablet and the VR condition, it should not have favored any of the hypotheses.

### Setup

The VR game was played with the HMD of the HTC Vive. A high-end computer (GForce GTX 780 Ti, Intel Core i7 4930, 16GB RAM) that was capable of rendering the game on more than 90fps was used. In the tablet condition, the game was played on a Samsung Galaxy Tab S3 tablet with Android Nougat 7.0. The text the subjects had to read in the control condition was also presented on the Galaxy Tab S3. To increase the level of immersion in the VR condition, the Bose QuietComfort 15 noise-canceling headphones were used. As the speakers of the Samsung Galaxy Tab S3 have a lower sound quality than the Bose Headphones, the UE Boom by Logitech was used to play the sound in the tablet condition to decrease the differences in the level of sound quality. Both conditions were played with the same loudness of about 50dB.

For the text of the control condition, the font Times New Roman was chosen as it is often used and well-known, which ensured a pleasant reading experience. Additionally, the serifs of the font increase the flow of reading. The line distance was set to 1.5 to make it easier for the participants to find the next line. The default font size was set to 11 pt. As participants were able to zoom in on the tablet, they could set their preferred font size themselves.

Participants could choose to be tested at either the University Medical Center Hamburg-Eppendorf or the informatics department of the University of Hamburg. They were not allowed to switch between the two locations. The testing rooms of both institutes were of average size (about 14m^2^) and contained few tables, chairs, computers and the camera stations of the VR. The rooms were quiet and distraction free. The participants were seated in a revolving chair throughout the whole experiment. The questionnaires were implemented using Google Forms and were presented to the participant on the computer. The brightness of the screens was kept constant throughout the whole experiment. The setup for both the VR and the tablet condition is shown in figure 2.

**Figure 2.**
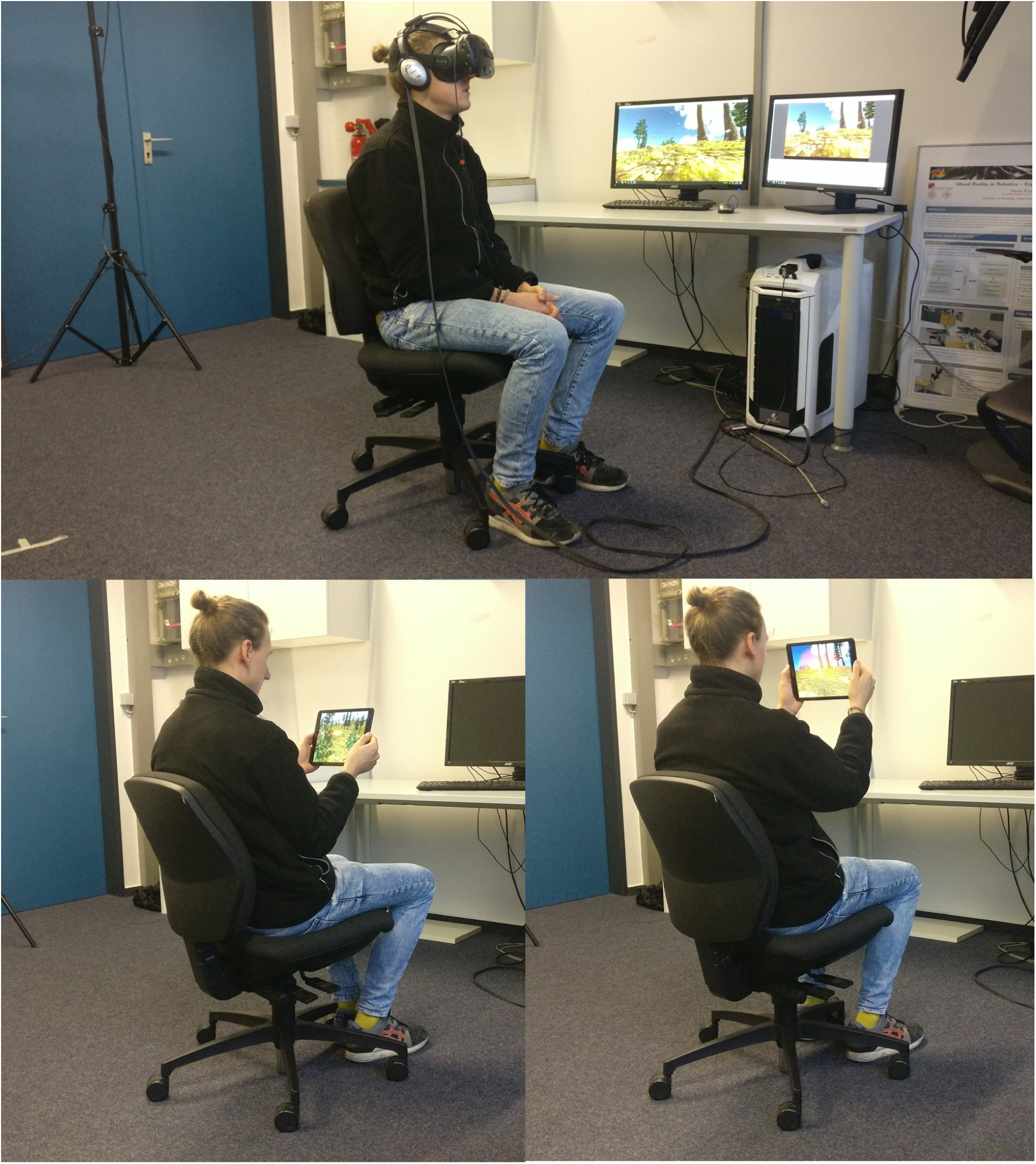
The setup of both the VR (top) and the tablet (bottom) condition. In the upper picture the player is sitting on a revolving chair wearing the noise cancelling headphones and the HMD of the HTC Vive. On the bottom left picture, the user is flying downwards and on the bottom right upwards. Sidewards rotation can be done with the upper body or chair.

**Figure 3.**
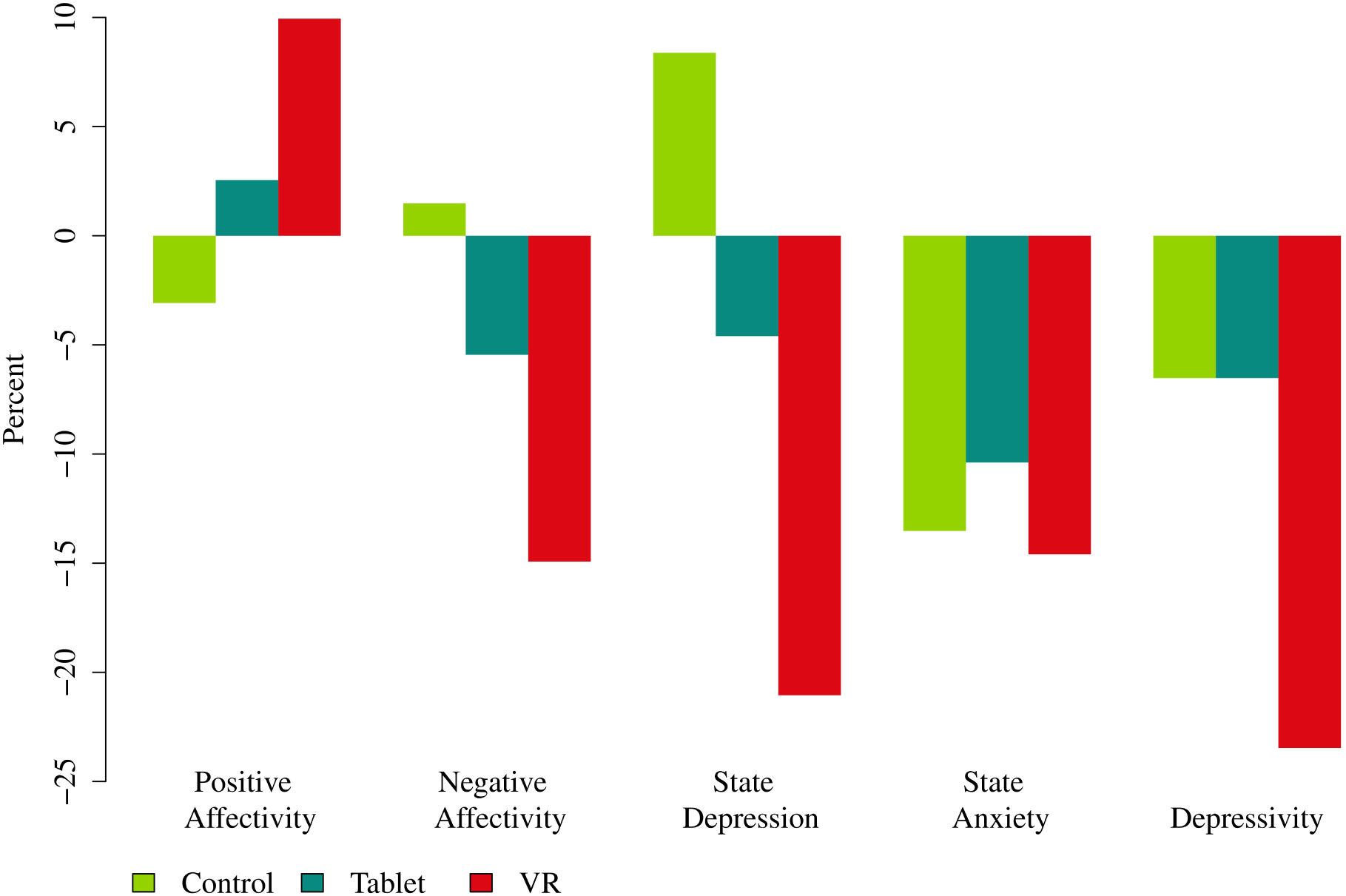
Percentage change of the primary affectivity variables between pre - and post-measures. Positive Affectivity and Negative Affectivity measured by the PANAS-X, State Depression and State Anxiety measured by the STADI(S) and Depressivity measured by the BDI-II (for more details see section Measures).

### Measures

The self-assessment questionnaires that were completed to measure the effects of different conditions on the subjects are explained below. The questionnaires were handed out before and after the condition, except for the last two (FSS and IPQ) which were only completed after each condition. All questionnaires were presented in the German version.

#### Positive Affect Negative Affect Schedule-Expanded Form (PANAS-X)

(Watson & Clark, 1999; Röcke & Grühn, 2003) The schedule includes 60 different adjectives of affect, whose intensity can be described on a ranging from 1=“very slightly or not at all” to 5=“extremely”. The participants were instructed to answer these questions in regard to their current affective state. The questionnaire was used to assess the subject’s current mood, through the variables Positive Affectivity and Negative Affectivity. Even though the term affectivity also refers to emotions, Positive Affectivity and Negative Affectivity are predominantly assessing mood (Watson & Clark, 1999). Emotional states are measured by the lower-order affects Fear, Sadness, Guilt, Hostility, Shyness, Fatigue, Surprise, Joviality, Self-assurance, Attentiveness and Serenity. Studies report high internal validity and good test-retest reliability (Watson & Clark, 1999).

#### State-Trait-Anxiety-Depression-Inventory (STADI)

(Laux, Hock, Ralf, Volker, & Karl-Heinz, 2013) The inventory includes a state and a trait questionnaire, each containing 20 affective statements. Subjects can indicate how much they agree with each statement on a ranging from 1=“not at all” to 4=“very much”. Both questionnaires assess the affects Euthymia, Dysthymia, Emotionality and Worry with 5 questions each. These are used to assess the intensity of the higher-order affects Depression, described by Dysthymia and Anhedonia (inverted values of Euthymia) and Anxiety, described by Emotionality and Worry. The state questionnaire is used to measures current feelings of depression and anxiety whilst the trait questionnaire assesses them as more long-living personality traits. This relatively new inventory has demonstrated a good reliability and validity (Brähler, Schumacher, & Strauβ, 2002).

#### Becks Depression Inventory II (BDI-II)

(Hautzinger, Keller, & Kühner, 2006) The inventory includes 2f questions regarding symptoms of depression over the past two weeks. Each question has 4 answers which are ordered in their intensity and evaluated on a ranging from 0 to 3. Higher scores indicate more intense symptoms. The BDI-II has a high internal consistency, a decent test-retest reliability for non-clinical samples and is sensitive to changes (Kühner, Bürger, Keller, & Hautzinger, 2007).

#### Simulator Sickness Questionnaire (SSQ)

(Kennedy, Lane, Berbaum, & Lilienthal, 1993) A questionnaire for the measurement of simulator sickness. Subjects can indicate the intensity of 16 symptoms regarding simulator sickness on a ranging from 0=“none” to 3=“severe”. The questionnaire is a standard questionnaire to measure simulator sickness and its validity has been shown (Kennedy et al., 1993).

#### iGroup Presence Questionnaire (IPQ)

(Schubert, Friedmann, & Regenbrecht, 2001) The questionnaire contains 14 items that assess the sense of presence including the sub-scales Spatial Presence, Involvement, Experienced Realism and the Sense of Being There. Subjects can indicate how much they agree with each statement on a ranging from 0 to 6. The questionnaire has demonstrated a good internal consistency (Schubert, Friedmann, & Regenbrecht, 1999).

#### Flow Short Scale (FSS)

(Rheinberg, Engeser, & Vollmeyer, 2002) The scale includes seven questions regarding components of the flow-experience which can be expressed on a ranging from 0=“does not apply” to 6=“does apply”. Three questions assess the worry that one could have about failing at accomplishing the task. In addition, three 9-point scales evaluate the self-estimated skill level, the self-estimated difficulty level of the task and the self-estimated feeling of overtaxing. The scale has been validated and shows a good internal consistency (Rheinberg, Vollmeyer, & Engeser, 2003; Engeser & Rheinberg, 2008).

### VR and Tablet Game

The game FlowVR was developed with the game-engine Unity, Version 5.6.3p1 for both HMD of the HTC Vive and an Android tablet. As nature has often been proven to induce positive feelings, reduce stress and improve mental health (Bratman, Hamilton, & Daily, 2012; Barton & Pretty, 2010), a grassland scene was chosen for the game. Grass, bushes, trees and terrain textures were acquired through the Landscape Auto Material Asset from the Unity Asset Store^6^. This setting is also beneficial since it contains mainly blue and green colors, which are associated with pleasure and neutral levels of arousal (Valdez & Mehrabian, 1994). The surface of the terrain contains various smooth and gentle hills that occur in a low to medium frequency (see figure 1), which is associated with gentle and playful moods (Poffenberger & Barrows, 1924). The flowers are ordered in such way, that they altogether build a path through the terrain. To highlight them visually and add variety to the scene, the flowers were given different, bright colors (blue, red, orange, yellow, purple and white) and were surrounded by a halo of the same color (see figure 1). Throughout the game, little leaves start circling around the user’s view (see figure 1). They indicate the flowers that have already been pollinated and thus represent another type of positive feedback to support a feeling of flow.

To find the right soundtrack for the game, three songs were selected by the criteria mentioned in the introduction and evaluated in a pilot study. 9 Participants (mean=30, sd=14.22, 5 female) had to listen to each song on three different days at around the same time of day on their phone and fill out the STADI(S)^7^ before and after. The song Silver Blue Light by Kevin MacLeod^8^ resulted in the most improvement of anxiety and depression (measured by the STADI(S)) and thus was implemented into the game.

### Procedure

On the first test-day participants were handed out the information and consent form. The subjects were informed that the study was interested in the subject’s feelings when playing the game and they were provided with a summary of the experiments procedure. At the end of the last test day, the remuneration of 25 Euros/class credits was paid and detailed information about the study’s purpose was provided.

After entering the room and sitting down, the participants had to fill out the STADI(S), PANAS-X, BDI and SSQ questionnaires on a computer. Thereafter, they had to either read a glider flight report (control condition), play FlowVR on a tablet (tablet condition) or play FlowVR in VR with a HMD (VR condition). The order of conditions was counterbalanced across participants.

In the control condition, participants had 5:50 minutes to read the flight report, which is exactly the time they would play the game in the VR and the tablet condition. Participants were told there is a given time to read the text, but that it is not important to finish the text by then, so they would not have to hurry.

Before playing the game in the VR and tablet condition, the participants were given information about the gameplay. They were asked to imagine to be a bee, whose goal it is to make the landscape flourish by flying above it and touch flowers or trees which are surrounded by a bright halo. That would lead to pollen popping out of the flowers and new flowers, bushes or trees growing all around. They were told to follow the path of the flowers and collect as many flowers as possible, however, they should not mind when missing out a flower. To make sure the participants would not struggle to control the game, the experimenter gave a short demonstration of the controls. In the VR condition, this was simply done by showing how to rotate the head when they wanted to fly in a certain direction (see figure 2). As the controls of the tablet are a little less intuitive, even though they followed the same logic as those of the VR, the instruction in the tablet condition was more detailed. Participants were told to imagine an arrow that sticks out of the back of the tablet, which indicates the flying direction. Therefore, they would have to rotate the back of the tablet toward the direction they wanted to fly. This could be done by either using their wrists or by moving the tablet within a spherical radius around them. To make sure their arm muscles would not be strained, they were able to place their arms on the armrest or their upper body, except when they were flying upwards (see figure 2). In both the VR and the tablet conditions they were allowed to use the revolving chair for the sideways rotation.

After 5:50 minutes both the reading and the gaming time were over. This time was set, as the song of the game lasted exactly that long and as the exposure to nature of about 5 minutes has been shown to result in a rather optimal mood improvement (Barton & Pretty, 2010). Furthermore, prolonged exposure to VR may have increased the risk of simulator sickness (Kolasinski, 1995).

In the game, participants would automatically stop flying and a text would show up, asking them to contact the experimenter. By then, the song was over and when the subjects followed the path, they also reached the last flower due to the continuous flying-speed. In the reading condition, the experimenter instructed them to finish their current sentence and stop reading. Finally, again the STADI(S), PANAS-X, BDI and SSQ were filled out. Additionally, the FSS and the IPQ were completed. When filling out the IPQ for the reading condition, the participants were asked to answer the questions in regard to how well they could delve into the story of the flight report. After completing the last questionnaire, the experiment day was over.

## Statistical Analyses

The primary affectivity variables are Positive Affectivity and Negative Affectivity measured by the PANAS-X, State Depression and State Anxiety measured by the STADI(S) and Depressivity measured by the BDI-II. Additionally, Simulator Sickness was measured by the SSQ, Sense of Presence was measured by the IPQ and Flow was measured by the FSS.

First, the conditions were examined independently of each other to test for significant changes in those variables that were measured before and after the conditions. Many distributions of the data were not normally distributed according to the Shapiro-Wilk-Test (see table 1). Consequently, the Wilcoxon-Mann-Whitney-Test was used. The results of all statistical tests are shown in table 4.

**Table 1.** P-values of the Shapiro-Wilk test for normal distribution.

| Variables | Control |  |  | Tablet |  |  | VR |  |  |
| --- | --- | --- | --- | --- | --- | --- | --- | --- | --- |
| | Pre | Post | $\Delta$ | Pre | Post | $\Delta$ | Pre | Post | $\Delta$ |
| BDI-II |  |  |  |  |  |  |  |  |  |
| Depressivity | **0.004** | **0.001** | 0.586 | **0.006** | 0.255 | 0.076 | 0.486 | 0.300 | 0.335 |
| STADI(S) |  |  |  |  |  |  |  |  |  |
| State Depression | **0.001** | 0.255 | 0.549 | **0.000** | **0.005** | 0.595 | 0.206 | **0.001** | **0.021** |
| State Anxiety | **0.001** | **0.000** | 0.484 | **0.004** | 0.273 | **0.020** | 0.931 | 0.097 | 0.306 |
| PANAS-X |  |  |  |  |  |  |  |  |  |
| Negative Affectivity | 0.189 | **0.003** | 0.131 | 0.590 | 0.113 | 0.098 | **0.004** | 0.243 | **0.000** |
| Positive Affectivity | **0.022** | **0.000** | 0.535 | **0.002** | 0.577 | 0.853 | **0.006** | 0.987 | 0.088 |
| SSQ |  |  |  |  |  |  |  |  |  |
| Simulator Sickness | **0.001** | 0.153 | 0.063 | **0.000** | 0.171 | **0.015** | **0.001** | **0.022** | 0.531 |
| IPQ |  |  |  |  |  |  |  |  |  |
| Sense of Presence |  | 0.254 |  |  | 0.084 |  |  | 0.249 |  |
| FSS |  |  |  |  |  |  |  |  |  |
| Flow |  | 0.693 |  |  | 0.079 |  |  | 0.912 |  |

*Note.* P-values below 0.05 are written in bold and indicate a non-normal distribution.
$\Delta$ = Difference between pre and post values.

The Friedman test was then applied to the data to test for significant differences between all three conditions. The Friedman-Test was chosen instead of a repeated measures ANOVA due to the non-normal distribution of some of the data. Of the variables that were measured before and after the conditions, the differences were calculated to use them as dependent variables for comparing the conditions.

To compare the conditions one by one, a pairwise Wilcoxon-Mann-Whitney-Test was then applied to the variables that were significant in the Friedman-Test. As the choice of the post-hoc test has an impact on the results, the tests were carried out with both the Bonferroni (BF) and the Benjamini & Hochberg (BH) procedure. Both are controlling for type 1 errors, which occur when the null hypothesis is falsely rejected. The BF method is known to be more stringent in correcting these errors, but on the flip-side, has a higher risk of failing to reject the null hypothesis even if it is actually false (type 2 error) (Shaffer, 1995). The resulting p-values are shown in table 5. Significance was set at an alpha of 0.05.

## Results

### Descriptive Statistic

All 18 participants managed to control the game and were able to follow the path of the flowers in both the VR and the tablet condition. They had an average BDI-II score of 6.67 (*min* = 4, *max* = 17, *sd* = 3.36) when applying for the study. The BDI-II mean was lower when assessed by the BDI-II questionnaires given prior to every condition during the actual experiment, however, this difference was not significant (*p* = 0.07). Throughout all three conditions, the pre-mean of the BDI-II was at 5.22 (*min* = 0, *max* = 17, *sd* = 4.26), with some participants having a BDI-II score lower than the targeted value of at least 4. As assessed by the STADI(T), the mean value of Trait Depression was at 11 (*min* = 5, *max* = 18, *sd* = 3.86) and the mean value of Trait Anxiety was at 15 (*min* = 10, *max* = 29, *sd* = 5.07). Means and standard deviations of pre and post measures can be found in table 2. The percentage change of the primary affectivity variables is presented in the bar chart 3. The means and standard deviations of these changes and of the ratings of Sense of Presence, Flow and Simulator Sickness are shown in table 3.

**Table 2.**
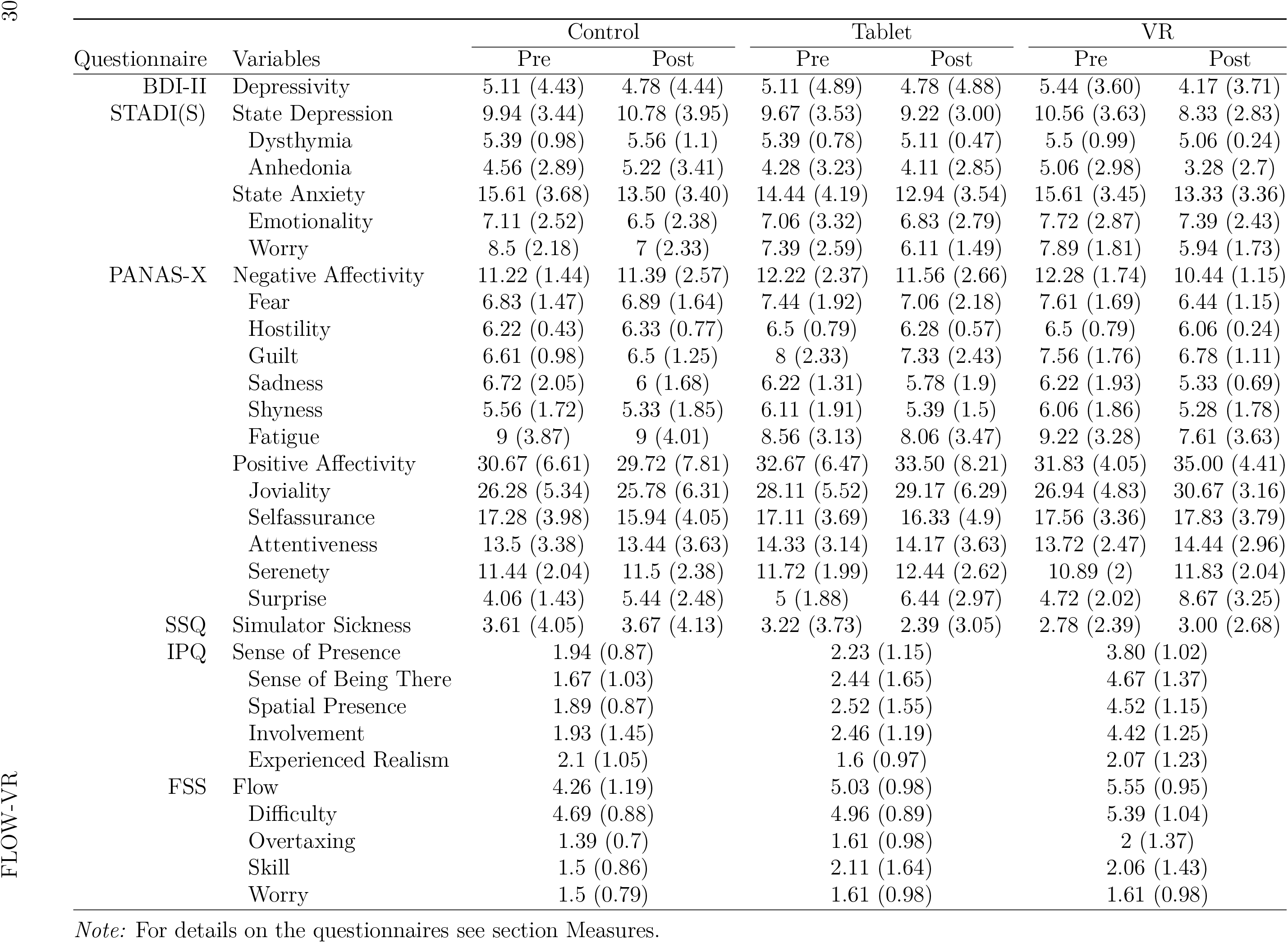
Means (and standard deviations) of all outcome variables.

**Table 3.** Means (standard deviations) and p-values of the pre-post differences of the primary ajjectivity variables and Simulator Sickness, plus the ratings of Flow and Sense of Presence.

| Variable | Control |  | Tablet |  | VR |  |
| --- | --- | --- | --- | --- | --- | --- |
|  | *M (SD)* | *p* | *M (SD)* | *p* | *M (SD)* | *p* |
| BDI-II |  |  |  |  |  |  |
| Depressivity | -0.33 (0.91) | 0.140 | -0.33 (0.91) | 0.193 | -1.28 (2.02) | 0.018* |
| STADI(S) |  |  |  |  |  |  |
| State Depression | 0.83 (1.89) | 0.133 | -0.44 (1.82) | 0.338 | -2.22 (2.51) | 0.004** |
| State Anxiety | -2.11 (2.54) | 0.005* | -1.5 (2.31) | 0.022* | -2.28 (3.34) | 0.014* |
| PANAS-X |  |  |  |  |  |  |
| Negative Affectivity | 0.17 (2.01) | 0.856 | -0.67 (1.57) | 0.134 | -1.83 (1.47) | 0.001** |
| Positive Affectivity | -0.94 (4.5) | 0.444 | 0.83 (4.06) | 0.450 | 3.17 (3.78) | 0.005* |
| SSQ |  |  |  |  |  |  |
| Simulator Sickness | 0.06 (1.21) | 0.745 | -0.83 (1.47) | 0.046* | 0.22 (2.8) | 0.861 |
| FSS |  |  |  |  |  |  |
| Flow | 4.26 (1.19) |  | 5.03 (0.98) |  | 5.55 (0.95) |  |
| IPQ |  |  |  |  |  |  |
| Sense of Presence | 1.94 (0.87) |  | 2.23 (1.15) |  | 3.8 (1.02) |  |

*Note:* For details on the questionnaires see section Measures.
\* $p < 0.05$ . \*\* $p < 0.005$ . \*\*\* $p < 0.0005$ .

**Table 4.** P-values of the pre-post-differences of all variables.

| Questionnaire | Variables | Control | Tablet | VR |
| --- | --- | --- | --- | --- |
| BDI-II | Depressivity | 0.140 | 0.193 | 0.018* |
| STADI(S) | State Depression | 0.113 | 0.338 | 0.004** |
|  | Dysthymia | 0.345 | 0.174 | 0.054 |
|  | Anhedonia | 0.255 | 0.618 | 0.007** |
|  | State Anxiety | 0.005* | 0.022* | 0.014* |
|  | Emotionality | 0.185 | 0.457 | 0.546 |
|  | Worry | 0.012* | 0.013* | 0.004** |
| PANAS-X | Negative Affectivity | 0.856 | 0.134 | 0.001** |
|  | Fear | 1.000 | 0.205 | 0.003** |
|  | Hostility | 0.572 | 0.330 | 0.026* |
|  | Guilt | 0.795 | 0.089 | 0.069 |
|  | Sadness | 0.151 | 0.179 | 0.037* |
|  | Shyness | 0.407 | 0.081 | 0.114 |
|  | Fatigue | 0.942 | 0.163 | 0.022* |
|  | Positive Affectivity | 0.444 | 0.450 | 0.005* |
|  | Joviality | 0.406 | 0.276 | 0.004** |
|  | Self-assurance | 0.037* | 0.146 | 0.694 |
|  | Attentiveness | 0.642 | 0.494 | 0.069 |
|  | Serenity | 1.000 | 0.225 | 0.089 |
|  | Surprise | 0.014* | 0.035* | 0.002** |
| SSQ | Simulator Sickness | 0.745 | 0.046* | 0.861 |

*Note:* For details on the questionnaires see section Measures.
\* $p < 0.05$ . \*\* $p < 0.005$ . \*\*\* $p < 0.0005$ .

It is noticeable, that VR has the highest values in reducing Depressivity, State Depression, State Anxiety and Negative Affectivity and in increasing Positive Affectivity. Furthermore, the highest values of Flow and Sense of Presence were also produced by the VR condition. For each of these variables, the VR condition is followed by the tablet condition with the second highest values. The control condition produced the lowest values.

Consequently, any significant difference between the conditions supports the assumption, that the VR condition is most effective in improving overall affectivity, followed by the tablet condition and lastly the control condition. Except for Simulator Sickness, where the tablet condition produced the highest reduction, followed by the control condition and lastly the VR condition.

### Inferential Statistics

#### Affectivity

The control condition and the tablet condition led to significantly decreased values of State Anxiety (control: *p* = 0.005^**^/tablet: *p* = 0.022^*^), with the main reason appearing to be the decline of the lower-order factor Worry (control: *p* = 0.012^**^/tablet: *p* = 0.013^*^) as Emotionality (control: *p* = 0.185/tablet: *p* = 0.47) was not significantly reduced. The other primary affectivity variables were not found to have significantly changed in both conditions.

The VR-condition led to significantly improved values in Depressivity (*p* = 0.018*), Positive Affectivity (*p* = 0.005 * *), Negative Affectivity (*p* = 0.0006^***^), State Depression (*p* = 0.004^**^) and State Anxiety (*p* = 0.014^*^). The decrease of State Depression is predominantly attributable to the decrease of Anhedonia (*p* = 0.007^**^), since Dysthymia was not reduced significantly (*p* = 0.054). Similarly, the decrease of State Anxiety is predominantly attributable to the decrease of Worry (*p* = 0.0036^**^), since Emotionality was not reduced significantly (*p* = 0.55). With these results, the null-hypothesis of the first hypothesis (H1) cannot be rejected since the only the VR game led to a significant improvement of mood and depression.

Significant differences in the Friedman-Test were found for State Depression (*χ*^2^ = 18.98,*p* = 0.0001^***^), Negative Affectivity (*χ*^2^ = 11.43, *p* = 0.003^**^) and Positive Affectivity (*χ*^2^ = 6.79, *p* = 0.034*). Depressivity (*χ*^2^ = 4.84, *p* = 0.089) and State Anxiety (*χ*^2^ = 1.16, *p* = 0.56) did not show significant effects. Thus, only State Depression, Negative Affectivity and Positive Affectivity were used for a pairwise comparison of the conditions.

When compared to the control condition, the VR condition produced significantly higher values of pre-post-difference in State Depression (BH/BF: *p* = 0.002^**^, *r* = 0.6), Positive Affectivity (BH/BF: *p* = 0.001^**^, *r* = 0.48) and Negative Affectivity (BH/BF: *p* = 0.001^**^, *r* = 0.49). No significant differences in the affectivity variables could be found between the tablet and the control condition. The choice of the post-hoc method is especially having an impact on the results when comparing the VR and the tablet condition. State Depression (BH: *p* = 0.023*/BF: *p* = 0.046*, *r* = 0.39) had significantly different values with both post-hoc tests. However, differences of Positive Affectivity (BH: *p* = 0.04*/BF: *p* = 0.079, *r* = 0.4) and Negative Affectivity (BH: *p* = 0.047*/BF: *p* = 0.095, *r* = 0.38) are only significant when using the BH multiple test correction.

As a significant difference between the tablet and the VR condition could be found regardless of the post-hoc method chosen, the depression-part of the third hypothesis is supported by the results. As Positive Affectivity and Negative Affectivity were significantly different when using the BH method but not when using the BF method, no conclusive statement can be made about whether there is an actual impact of immersion on mood enhancement as stated by the third hypothesis (H3).

The changes of the lower-order affects of the PANAS-X, namely Fear (*p* = 0.117), Guilt (*p* = 0.192), Sadness (*p* = 0.607), Joviality (*p* = 0.077), Self-assurance (*p* = 0.168), Attentiveness (*p* = 0.238), Shyness (*p* = 0.984), Fatigue (*p* = 0.068), Serenity (*p* = 0.130) and Surprise (*p* = 0.083) were not significantly different according to the Friedman test. Except for Hostility (*p* = 0.039*), which was not significant in the subsequent pairwise Wilcoxon-Mann-Whitney-Test (see table 5).

**Table 5.** Controlled p-values of the pairwise comparison of all variables that produced significance in the Friedman test.

| Variables | Control<>Tablet |  | Control<>VR |  | VR<>Tablet |  |
| --- | --- | --- | --- | --- | --- | --- |
|  | BH<sup>a</sup> | BF<sup>b</sup> | BH | BF | BH | BF |
| BDI-II |  |  |  |  |  |  |
| State Depression | 0.077 | 0.230 | 0.002** | 0.002** | 0.023* | 0.046* |
| Anhedonia | 0.175 | 0.526 | 0.005 | 0.005 | 0.014 | 0.027 |
| PANAS-X |  |  |  |  |  |  |
| Negative Affectivity | 0.138 | 0.415 | 0.013* | 0.013* | 0.047* | 0.095 |
| Hostility | 0.296 | 0.592 | 0.155 | 0.155 | 0.491 | 1.000 |
| Positive Affectivity | 0.275 | 0.824 | 0.012* | 0.012* | 0.04* | 0.079 |
| FSS |  |  |  |  |  |  |
| Flow | 0.043* | 0.087 | 0.026* | 0.026* | 0.088 | 0.265 |
| Worry | 0.200 | 0.600 | 0.099 | 0.099 | 0.200 | 0.460 |
| Difficulty | 0.551 | 1.000 | 0.336 | 0.336 | 0.551 | 1.000 |
| IPQ |  |  |  |  |  |  |
| Sense of Presence | 0.433 | 1.000 | 0.002** | 0.004** | 0.002** | 0.004** |
| Sense of Being There | 0.158 | 0.475 | 0.002** | 0.002** | 0.002** | 0.003** |
| Spatial Presence | 0.124 | 0.371 | 0.001** | 0.001** | 0.002** | 0.004** |
| Involvement | 0.316 | 0.948 | 0.002** | 0.004** | 0.002** | 0.002** |

*Note:* For details on the questionnaires see section Measures.
\* $p < 0.05$ . \*\* $p < 0.005$ . \*\*\* $p < 0.0005$ . <sup>a</sup> BF = Bonferroni post-hoc method.
<sup>b</sup> BH = Benjamini & Hochberg post-hoc method.

Anhedonia was the only lower-order effect of the STADI(S) that was found to have significant pre-post-differences between the conditions (*χ*^2^ = 18.72,*p* = 0.0001^***^). According to the pairwise Wilcoxon-Mann-Whitney-Test both VR and control (BH/BF: *p* = 0.005^**^, *r* = 0.57) and VR and tablet (BH: *p* = 0.014*/BF: *p* = 0.027*, *r* = 0.44) were significantly different. Dysthymia (*p* = 0.072), Emotionality (*p* = 0.459) and Worry ((*p* = 0.410) did not change significantly different between the conditions.

#### Simulator Sickness

The tablet condition was the only condition that significantly reduced Simulator Sickness (*p* = 0.046*). Both the control and the VR condition did not significantly impact Simulator Sickness. Changes of Simulator Sickness were not significantly different between the conditions (*χ*^2^ = 3.6,*p* = 0.17). Consequently, Simulator Sickness was successfully avoided in all conditions, therefore should not have had any negative impact on the results.

#### Sense of Presence

As expected, values of Sense of Presence were found to be significantly different between the conditions (*χ*^2^ = 21.33,*p* = 0.0001***). The VR condition produced significantly higher values compared to both the tablet (BH: *p* = 0.002**/BF: *p* = 0.004**, *r* = 0.54) and the control condition (BH: *p* = 0.002**/BF: *p* = 0.004**, *r* = 0.54). These results support the second hypothesis (H2). The tablet and the control conditions were not significantly different (BH: *p* = 0.433/BF: *p* = 1), which means participants were about equally good in imagining the story and delving into the non-immersive game. No significant correlation between Sense of Presence and State Depression has been found in any of the conditions (VR: *p* = 0.074, Tablet: *p* = 0.077, Control: *p* = 0.26).

#### Flow

Levels of Flow were also significantly different between the conditions (*χ*^2^ = 8.82,*p* = 0.012*). The VR and the control condition were significantly different (BH: *p* = 0.026*/BF: *p* = 0.026*, *r* = 0.4). A significant difference between the tablet and the control condition was found only when using the BH procedure (BH: *p* = 0.043*/BF: *p* = 0.087, *r* = 0.36). Tablet and VR were not significantly different. Additionally, Flow was negatively correlated with State Depression in both the tablet (*p* = 0.007, *r* = —0.61) and the VR (*p* = 0.013, *r* = —0.57) condition.

Self-estimated skill (*p* = 0.07) and feelings of overtaxing (*p* = 0.88) measured by the FSS were not significantly different between all three conditions, whilst worry (*χ*^2^ = 6,*p* = 0.0498*) and self-estimated task-difficulty (*χ*^2^ = 8.07,*p* = 0.018*) were. However, the subsequent pairwise comparison applied for the latter two did not reveal any significant difference between any of the conditions (see table 5). Since the balance between difficulty and skill seems to be quite similar in all three conditions, there should be no excessive difference between the conditions in feelings of boredom or overtaxing.

## Discussion

Former research has successfully utilized VR for different types of anxiety disorders. However, a recent meta-analysis revealed, that VR therapy for other mental disorders, such as depression, has rarely been researched (Freeman et al., 2017). To investigate the question of whether VR can be efficiently used for improving depressive mood, the game FlowVR was developed and designed to improve mood and reduce feelings of depression. Subsequently, FlowVR’s impact on affectivity was tested in VR and on a tablet. The evaluation showed, that the VR game was successful in improving overall affectivity and temporarily reducing depression.

The VR condition led to significant improvements in all questionnaires regarding affectivity and produced significantly better improvements in variables that assessed mood and depression compared to the control condition. This finding indicates that VR can be efficiently utilized to enhance depressive mood. Apart from State Anxiety, the tablet and the control conditions did not improve any dependent variable regarding affectivity, such as State Depression, Negative Affectivity or Positive Affectivity. Furthermore, no significant difference could be found between the tablet and control condition in any of the questionnaires. Thus, playing FlowVR on a tablet does not seem to offer any advantages compared to reading the text. As expected, the VR condition had significantly higher values in Sense of Presence when compared to the tablet and the control condition, which is likely caused by its higher level of immersion. Thus, the positive elements implemented into the game do not seem to be sufficient to actually improve depressive mood but only seems to work in combination with a higher sense of presence.

In comparison to the tablet condition, the VR condition had significantly better results in the reduction of State Depression. The significant result of State Depression can mainly be attributed to the improvement of the values in Anhedonia, which is determined by measuring Euthymia and inverting its values. Since Euthymia is a positive affect, the game seems to have been particularly successful in improving depression by inducing positive affects. If the BH post-hoc test is chosen, the VR condition also led to significantly higher increases in Positive Affectivity and reductions in Negative Affectivity, with medium to high effect sizes. Additionally, whilst the VR condition was significant in every affect-questionnaire, the tablet condition only produced significant improvement in State Anxiety. Bearing the above in mind, the VR seems to be more effective for mood improvement than the tablet. Since State Depression was strongly negatively correlated with Flow but not with Presence in both the VR and the tablet condition, the games positive impact on depression seems to be predominantly related to the experience of flow.

As all conditions have similarly decreased values in State Anxiety, factors other than differences between the conditions might have been the reason for this phenomenon. The conditions led to significant decreases in Worry, however not in Emotionality. Thus, the decrease could be due to a reduction in test anxiety. Regardless of the main influencing factor, neither the VR game nor the tablet game seems to be particularly more suitable for reducing anxiety than the control condition.

Interestingly, VR also significantly improved the values of Depressivity measured by the BDI-II, which assesses symptoms of depression over the last two weeks. This might be related to the mood-congruence effect, as the improved mood could have primed the evocation of memories and lead to a positively biased evaluation of the last weeks. However, differences in Depressivity between the VR and the control condition were not significant. Even if the VR condition might have affected the emotional evaluation of the past weeks, this effect does not appear to be excessively strong compared to the other conditions. Therefore, no statement can be made about the intensity of this effect.

### Limitations

Apart from the positive results, the study contained a few weaknesses which should be addressed in future studies. Even though the tablet and the VR conditions were designed to be as comparable as possible, some other factors than just the level of immersion could have interfered. Firstly, some participants might have been able to predict the hypothesis of the study. Participants were informed, that the study is interested in their feelings when playing the game. In combination with the experiment design, this might have revealed the study’s interest in improving affectivity with VR. This could have influenced participants to answer the mood questionnaires in favor of the hypothesis. However, one could argue that if that was the main reason for the different results, State Anxiety should also have been improved more in the VR condition than in the other conditions. It is rather unlikely, that participants have distorted their responses on depression but not on anxiety, as both mood disorders are related (Wittchen & Hoyer, 2011) and the study did not reveal its particular interest in depression.

Secondly, as the sound was played from different devices, the different sound quality could have also had an impact. The qualities of the speaker and the headphone were broadly comparable but still, the headphones quality was higher. The decision to use different devices was made, as noise-canceling headphones are highly immersive from the auditory perspective. They filter out ambient noises so that hardly any external auditory stimuli can disturb the listener. As this study is interested in the impact of immersion, the tablet condition had to have a non-immersive sound in order to serve as a non-immersive comparison.

The controls followed the same logic, however, some participants found the controls of the tablet a little more difficult than those of the VR. Due to the different nature of the devices, these discrepancies could hardly be avoided. In fact, the higher level of immersion in the VR condition might have been a reason for reporting easier and more intuitive controls. The resulting increase in difficulty might have made the subjects feel overtaxed and left a negative impact on their mood. However, the self-estimated difficulty of the conditions and self-estimated feelings of overtaxing, both measured by the FSS, were not significantly different between the conditions. Additionally, State Anxiety similarly decreased in both conditions, which contradicts a feeling of overtaxing in the tablet condition. Furthermore, all participants managed to control the game and follow the path of the flowers with both devices. Because subjects had to hold the tablet in their hands, this burden on the arms and hands could have also had an impact. However, they were able to rest their arms on the upper body if not flying upwards and no participant complained to have been overly strained. Lastly, since most of the participants were under 30 years old and of Western European origin, it should be noted that the results cannot necessarily be generalized for older people or those of other ethnicities.

### Future Work

As the game only tested short-term effects with healthy participants, much more research has to be conducted to ascertain whether VR can efficiently be utilized for depression therapy. The effects of using the game on a daily or weekly basis should be investigated in the future. Participants might be able to keep their elevated mood or improve it throughout the time. In this case, the hazard of becoming addicted should also be investigated. However, they could also get used to the game and the positive effect might disappear. In order to avoid this, the game could be extended, with different levels to keep up the subjects interest in the game.

Furthermore, subjects affected by depression might react differently to the VR game than healthy participants. Studies have shown that patients affected by a major depressive disorder are failing to show a positive bias when primed by a stimulus with a positive valence (LeMoult, Yoon, & Joormann, 2012). Participants might be less affected by the positive elements of the game and thus it might be more difficult to improve their mood. On the other hand, most of the participants that took part in this study were already in a rather good mood, with one of them even having a BDI-II score of 0. As it might be more difficult to improve the mood of participants that are already happy, a study with depressive subjects could also lead to better results in mood improvement.

Additionally, the game could be customized for the subjects. For example, the level of difficulty of the game could be balanced with the participant’s individual skill, to assure participants will experience flow. Moreover, the concepts of VR exposure therapy could be transferred to VR depression therapy. Objects that are related to the subject’s cause of depression could be embedded into the game. This could help patients to approach adverse memories and to experience them in a new context to eventually overcome them. Lastly, the game needs to be compared to other mood-improvement techniques, such as reading or imagining an emotional story. Some subjects might find it really difficult to delve into their imagination, thus should be more affected by VR.

### Conclusion

To conclude, the VR game was successful in improving depressive mood. A short exposure time of about six minutes led to an improved mood and reduced feelings of depression. Even when compared to the control and the tablet conditions, the results suggest that the VR condition was substantially more effective. However, whether the VR game is actually suitable and efficient for application in clinical contexts is not answered by this study. More research on its long-term effect has to be conducted and the VR game has to be tested with patients that are actually affected by depression. This study complements existing research, which to a certain extent has already been able to use VR to reduce depression in subjects. It substantiates that it is worthwhile to further invest in the research on VR depression therapy.

## Author Contributions

SK and JG conceived the presented idea. All authors conceived and planned the experiment. LB implemented FlowVR with the support from FM. LB carried out the experiment. LB analyzed the data and wrote the manuscript with support from FM and SK. All authors provided critical feedback and helped shape the research, analysis and manuscript.

## Data Availability Statement

The raw data supporting the conclusions of this manuscript will be made available by the authors, without undue reservation, to any qualified researcher.

## Footnotes

1 http://www.who.int/mediacentre/factsheets/fs369/en/ (accessed 21.06.2018).

2 https://cubicleninjas.com/portfolio/guided-meditation-vr.

3 http://www.exploredeep.com.

4 https://en.oxforddictionaries.com/definition/virtual_reality (accessed 21.06.2018).

5 www.vive.com(accessed 23.10.2018).

6 https://www.assetstore.unity3d.com

7 For more details on the STADI(S) see section.

8 http://incompetech.com/music/royalty-free/index.html?isrc=USUANl100718(accessed 21.06.2018).

